# Genome-wide association study reveals candidate loci for resistance to anthracnose in blueberry

**DOI:** 10.1101/2025.04.02.646913

**Authors:** Lushan Ghimire, Paul Adunola, Philip F. Harmon, Camila F. Azevedo, Tyler J. Schultz, Bruno Leme, Felix Enciso-Rodriguez, Juliana Benevenuto, Luis Felipe V. Ferrão, Patricio R. Munoz

**Author notes:** Correspondence: Corresponding Author: Patricio R. Munoz.

## Abstract

Anthracnose, caused by *Colletotrichum gloeosporioides*, poses a significant threat to blueberries, necessitating a deeper understanding of the genetic mechanisms underlying resistance to develop efficient breeding strategies. Here, we conducted a genome-wide association study on two distinct populations, comprising 355 advanced selections from the University of Florida Blueberry Breeding and Genomics Program. Visual scores and image analyses were used for assessing disease severity. The population was genotyped using Capture-Seq, detecting 38,379 single nucleotide polymorphisms. The study revealed a moderate narrow-sense heritability estimate (∼0.5) for anthracnose resistance in blueberries. Minor additive loci contributing to anthracnose resistance were identified on chromosomes 2, 3, 5, 6, 8, 9, 10, 11 and 12, spanning different populations. Image analyses demonstrated heightened sensitivity, detecting more associations within both populations compared to the visual approach. Candidate gene mining flanking significant associations unveiled key defense-related proteins, such as serine/threonine protein kinases, pentatricopeptide repeat-containing proteins, E3 ubiquitin ligases that have been well-known for their roles in plant defense signaling pathways. The dissection of speculated defense-related proteins into distinct layers offers further understanding into the intricate defense responses against *C. gloeosporioides*. Our findings highlight the complex and quantitative resistance mechanism for anthracnose in blueberry, providing insights for breeding strategies and sustainable disease management.

## Introduction

Anthracnose is a detrimental fungal disease of significant global importance, impacting a broad range of crops. The *Colletotrichum* genus, causing the disease, ranks among the top plant pathogen genera for its economic importance (Dean et al. 2012). Within it, *C. gloeosporioides* is reported to be the main cause of anthracnose in southern highbush blueberries (SHB, *Vaccinium corymbosum* and hybrids) (Velez-Climent and Harmon 2016; Phillips et al. 2018). It has also been reported to affect blueberries worldwide, in regions that include North America, Japan, Spain, South Korea, and China (Barrau et al. 2001; Yoshida and Tsukiboshi 2002; Polashock et al. 2005; Yu et al. 2005; Kim et al. 2009). Anthracnose-induced yield loss in blueberry has been documented to exceed 50% by the third harvesting cycle in New Jersey and under poor storage conditions, losses of up to 100% are possible (Polashock et al. 2017).

Anthracnose symptoms include sunken lesions with conidial masses, leading to substantial dieback, affecting various plant tissues, diminishing production, and rendering fruits unmarketable (Phillips et al. 2018). Conidia germinate and can initiate infection through direct penetration via appressoria or through wounds and natural openings (Jeffries et al. 1990). No commercial blueberry cultivars are immune to anthracnose; thus, fungicides remain the primary method to prevent crop loss. However, resistance to quinone outside inhibitor fungicides have already been reported in blueberry farms in Florida (Harmon 2014), hence the need for alternative methods of control for this disease.

Leveraging genetic sources of disease resistance presents a more sustainable and cost-effective alternative to chemical management (Polashock et al. 2005). However, improving host resistance through traditional breeding is time-consuming and resource intensive (Slater et al. 2013). Recent advancements in sequencing and automation technologies have substantially lowered the genotyping costs, facilitating the use of molecular markers to expedite trait selection. Disease resistance to anthracnose have been developed in several plant species such as grapes (Jang et al. 2020), pepper (Suwor et al. 2017), sorghum, (Ramasamy et al. 2009) and mango (Felipe et al. 2022) using modern strategies. In blueberries, genome-wide association studies (GWAS) have also been successfully implemented to elucidate the genetic architecture of various traits, such as fruit quality, off-season flowering, and flavor-related volatiles (Ferrão et al. 2018, 2020; da Silva et al. 2024).

Despite the economic importance of anthracnose in blueberries, the genetic basis of the disease resistance remains poorly understood. Most studies in blueberry have been conducted on anthracnose fruit rot (caused by *C. fioriniae*, formerly *C. acutatum*) during post-harvest stage (Polashock et al. 2005; Wharton and Schilder 2008; Miles et al. 2011; Miles and Hancock 2022; Jacobs et al. 2023). In this study, we evaluated the genetic architecture of anthracnose resistance across two southern highbush blueberry (SHB) populations for leaf and stem symptoms when inoculated with *C. gloeosporioides* using visual and image assessments. Potential variations in genetic architecture and resistance mechanisms might exist for these different pathogen species and blueberry populations used in previous studies (Osorio et al. 2014; Jacobs et al. 2019). Instead of fruit tissues, our assessment targeted leaf and stem tissue severity, aligning with findings indicating tissue-specific resistance mechanisms (Ehlenfeldt et al. 2006; Osorio et al. 2014; Jacobs et al. 2019). Understanding these intricacies in the genetic mechanisms is crucial for optimizing breeding strategies and enhancing anthracnose resistance. Hence, to our knowledge, our study represents the first investigation into the genetic architecture of resistance to anthracnose caused by *C. gloeosporioides* on leaves and stems of SHB blueberries. We explored the genetic and phenotypic variability in two breeding populations while comparing visual and computer vision phenomics approach, revealing marker associations and potential candidate genes underlying this trait. Our findings offer valuable insights into the genetic basis of resistance, informing strategic breeding decisions for anthracnose.

## Materials and methods

### Plant material

Phenotypic evaluations were performed in two SHB populations, consisting of advanced breeding selections from the University of Florida Blueberry Breeding Program. These genotypes were pre-selected for yield and fruit quality traits. Plant material for our experiment was collected from plants managed under commercial conditions in Waldo, FL, USA. The first population comprised 157 genotypes that were assessed for resistance in May 2023 (Popn-I). A second independent breeding population (Popn-II) comprised 198 genotypes and was assessed in August of 2023. To account for the between-experiment variation, we used 28 genotypes that were evaluated in the first population, as checks, in the second round of evaluation together with Popn-II. Six softwood cuttings each measuring approximately 40 cm in length, were collected from each genotype in both populations.

### Source of isolate and preparation of inoculum

A monoconidial isolate of *C. gloeosporioides* (isolate ‘15–646’) that was originally collected in 2015 from naturally infected stems of the ‘Flicker’ cultivar on a commercial blueberry farm in Central Florida was used to prepare inoculum for both experiments (Phillips et al. 2018). The isolate was grown on potato dextrose agar (PDA) incubated under continuous fluorescent lighting at 25 °C for 5 days. Conidia were quantified utilizing a hemacytometer (Bright-Line Hemacytometer; Hausser Scientific, Horsham, PA, USA) and the concentration of the suspension was adjusted to 1 × 10^7^ conidia mL^−1^ with autoclaved distilled water for spray application. The conidia suspension was diluted to 1 × 10^4^ conidia mL^−1^ for dip inoculations. Control plants were sprayed with equal volumes of distilled water.

### Inoculation procedure

Two softwood cuttings per genotype were randomly chosen and bundled for each replication (replicate-bundles), with a total of three replications per genotype each with a unique QR-code. These replicate-bundles were wounded on the lower 1/3^rd^ of the stem and placed over three 144.8 cm x 45.7 cm x 19.7 cm plastic troughs (Model: G5435768, Bayhead Products, Zoro; USA). Each trough represented a randomized complete block with one replicate (two cuttings) of each genotype. This entire experimental arrangement was housed within a self-constructed plastic tent situated in a growth room, consistently maintained at 90-95% relative humidity (Cool mist humidifier, Mikkin, USA) and 23°C temperature (Supplementary Fig 1A).

All the replicates were sprayed with the 1 × 10^7^ conidia mL^−1^ suspension until runoff, utilizing an aerosol spray gun (Crown Spra-Tool, Aervoe, NV, USA), thereby coating all stem surfaces and upper leaf surfaces. Each of the three troughs were filled with 40L of the 1 × 10^4^ conidia mL^−1^ suspension. The genotypes were monitored daily for 35 days after inoculation. Following the 35-day incubation period, infection by *C. gloeosporioides* was confirmed through isolations from necrotic lesion boundaries excised from five random genotypes selected from each block. Symptomatic tissue was surface sterilized with a 10% bleach solution for 1 minute, followed by three 1-minute water rinses. Tissue was plated onto PDA and incubated under continuous fluorescent light at 25 °C until conidia with characteristic morphologies consistent with the original reference isolate were observed.

### Disease phenotyping

In our study, we adopted two phenotyping approaches for disease quantification on day 35 post-inoculation: visual inspection of the individuals by human eyes (visual-based) and the use of a custom-made automatic computer vision phenotyping method (image-based). Visual evaluation involved scoring disease severity in a continuous scale, as a percentage of the total area covered by necrotic leaf and stem lesions. This score was recorded for each genotype within a replicate as an average value for the two stem cuttings per replicate. Simultaneously, comprehensive images encompassing all leaves and stem tissues for each genotype were captured, incorporating a unique QR-code. Subsequently, these images underwent disease assessment through the computer vision phenotyping approach.

The computer vision analysis employed a naive Bayesian machine learning methodology (Supplementary Fig 1B, C) (Ferrão et al., *in press*). A set of segmentation masks was generated based on a multi-dimensional statistical distribution, created through manual sampling of pixels in grid-samples (regions of features). These pixels were labeled under distinct feature classes, namely healthy, diseased, and background. To train the model, pixels were sampled extensively in 3×3 pixel grids from a diverse set of photos, ensuring the resulting probabilities reflected a representative population of the input data. The model, in this context, constitutes a probability density function (PDF) set containing probabilities for the appearance of features in the HSV colorspace. Subsequently, the disease severity percentage was calculated from segmentation mask areas as follows:

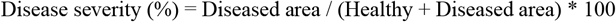

### Genotypic data

Fresh, juvenile leaves were harvested from each individual for subsequent genotyping procedures. DNA extraction and genotyping via Capture-Seq were performed at RAPiD Genomics (Gainesville, FL, USA). A genome-wide panel of 10,000 biotinylated probes, each spanning around 120-mers, were used for targeted capture, followed by high-throughput sequencing on the Illumina HiSeq2000 platform through 150-cycle paired-end runs.

SNP calling procedures adhered to the methodology by Ferrão et al. (2021). Briefly, raw reads underwent quality-based filtering and trimming. The filtered reads were aligned against the largest haploid scaffold set derived from the *V. corymbosum* cv. ‘Draper’ genome (Colle et al. 2019) using Mosaik v.2.2.3 (Lee et al. 2014). SNPs were subsequently called with the 10,000 probe positions as targets employing FreeBayes v.1.3.2 (Garrison and Marth 2012). SNP filtering criteria included a minimum mapping quality of 10, exclusivity to biallelic loci, a maximum missing data of 50% and a minor allele frequency (maf) threshold of 0.01. We used Vcftools v. 0.1.16 to extract sequencing read counts per allele and individuals from the variant call file (Danecek et al. 2011).

The updog package v.2.1.0 in R was employed to ascertain allele dosages based on the read counts (Gerard et al. 2018). SNPs accurately genotyped in 95% of the individuals were retained, utilizing the function ‘prop_mis’. The ploidy parameterization was encoded by the dosage of the alternative allele (B) in relation to the reference allele (A) as follows: 0 (AAAA), 1 (AAAB), 2 (AABB), 3 (ABBB), and 4 (BBBB) (Rosyara et al., 2016).

### Phenotypic analysis

We calculated the average of all three replicates within each population measured in the continuous scale. Also, in our study, we observed that some genotypes with moderate to severe anthracnose symptoms exhibited defoliation during incubation. In contrast, genotypes with lower symptom levels didn’t exhibit visible leaf drop (Fig1B). This observation raised concerns regarding the accuracy of severity assessment, particularly for genotypes falling within the moderate to severely symptomatic spectrum. To mitigate potential inaccuracies and improve precision in our analysis, we opted to simplify our dataset using a binary approach. We applied a stringent threshold of the 5^th^ percentile on the respective population’s average severity across three replicates. This categorization involved assigning highly resistant genotypes with no to minimal leaf drop to category 0, while grouping genotypes exhibiting moderate to severe symptoms, with moderate to high leaf drop incidence into category 1. By adopting this classification method, we aimed to reduce the likelihood of errors in severity assessment and focus on discerning the extremes ends of resistance phenotypes. For the analysis, in each of the scales (continuous and binary), the distinct datasets obtained from both the populations were combined (Comb_popn), and a one-stage correction was applied. This correction used the overlapping 27 genotypes from both populations as checks to correct for phenotyping batch effects and estimate the best linear unbiased estimates (BLUEs). Thus, adjusted means for each genotype were obtained using the following linear model:

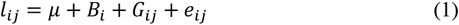

where *l*_*ij*_ is the liability for the phenotypic values represented by *y*_*ij*_, *y*_*ij*_ is the phenotype of genotype *i* in batch *j*; and they associated by the identity link function for the continuous scale and logit link function for binary scale; *μ* is the overall mean; *B*_*i*_ is the fixed batch effect; *G*_*ij*_ is the genotypic effect of individual *i* at batch *j*; and *e*_*ij*_ is the random residual effect. For the genotypic effect, the experiment was designed as an Augmented Block Design with common checks connecting different batches. Thus, the genotypic effect was separated into two groups, where *g* is the fixed effect for the regular individuals and *c* is the fixed effect associated to the checks. The BLUES of each genotype were used as the response variable in subsequent analyses. Additionally, we analyzed each population separately using both binary data (in binary scale) and the average values from three replicates (in continuous scale).

**Fig 1.**
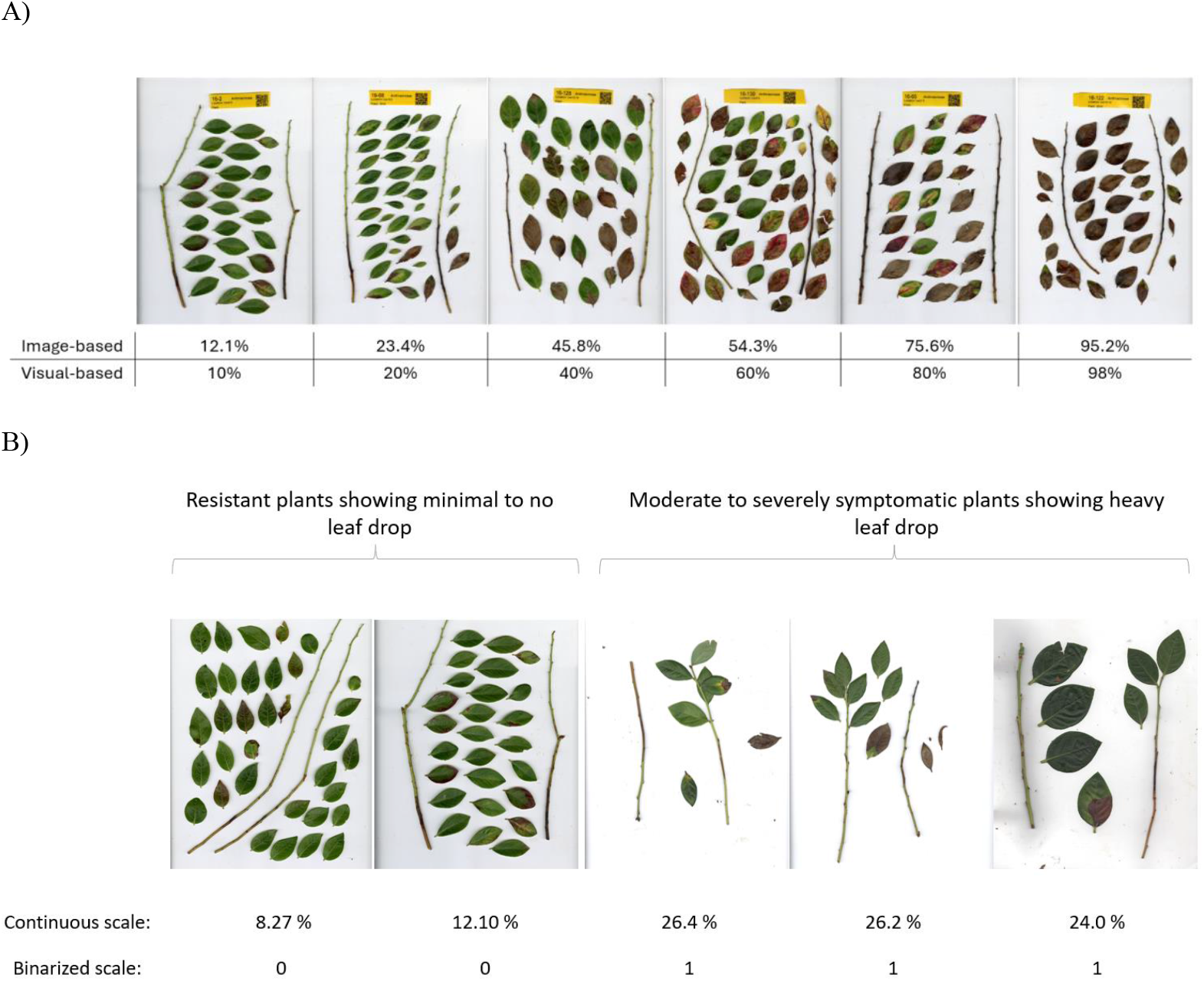
Severity assessment of different southern highbush blueberry genotypes for anthracnose and rationale for binarization. A) Severity scores of different genotypes in a continuous scale (disease severity percentage) in response to anthracnose using the two different phenotyping approaches: image- and visual-based approach. B) Comparative visualization of leaf drop in highly resistant versus moderately to severely symptomatic genotypes, showcasing severity scores on a continuous scale alongside binary categorization.

Furthermore, heritability estimates were computed to quantify the proportion of phenotypic variation attributed to genetic factors by considering genotypes as random effects and incorporating pedigree information into the relationship matrix: ***y***^**\***^ = **1***μ* + ***Za*** + ***e*** (2), where ***y***^**\***^ is the vector of BLUEs; μ represents the population mean with assumed flat prior distribution; ***e*** is the vector of random errors with 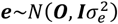 and 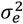 is the residual variance; and ***a*** is the vector of genomic estimated breeding values and its incidence matrix ***Z***, with 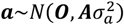, and 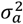 is the genetic variance. For variance components, we assumed a scaled inverted chi-square distribution. The construction of the pedigree relationship matrix (A) was done using the AGHmatrix package in R (Amadeu et al. 2016). This model was implemented by the BGLR package in the R software (Pérez and de los Campos 2014), defining 500,000 iterations for the Markov chain Monte Carlo (MCMC) algorithms, a burn-in period of 50,000 MCMC cycles, and thin equals to 10 before saving samples from each, totaling 45,000 MCMC cycles. The heritability (*h*^2^) was computed as the posterior mean: 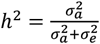

### Genome-wide association study

To identify the genomic regions regulating anthracnose resistance, GWAS was conducted using the R package GWASpoly v.2.11, specifically tailored for autopolyploid (Rosyara et al. 2016). For correction of population structure, the Q matrix was constructed using the top five principal components. Kinship matrices (K) were calculated using the algorithm embedded within the GWASpoly package. The testing of associations between SNPs and phenotypic variations was computed by the application of the Q + K linear mixed model (Rosyara et al. 2016). An additive genetic model where the SNP effect scaled proportionally to allele dosage was explored. Adjusted Bonferroni correction, considering a significance level of 0.05, was used for establishing a *p*-value detection threshold. Associations between SNPs and phenotypes were examined using quantile-quantile plots of estimated - log10(p) values.

### Candidate gene mining

For the exploration of potential candidate genes post-GWAS, we selected genomic regions within a 100 kb window upstream and downstream of significantly associated SNPs using the ‘Draper’ reference genome. The corresponding protein sequences were used as queries for comprehensive functional annotation using PANNZER (Törönen and Holm 2022) and eggNOG mapper (Huerta-Cepas et al. 2019). Candidate genes with functional description and gene ontology (GO) functions related to plant resistance or defense were selected and their potential functions were verified in literature search.

## Results

### Phenotypic analysis

Image- and visual-based assessments of disease severity after *C. gloeosporioides* inoculation were performed in two SHB populations (Popn-I and Popn-II). A significant correlation was observed between the two phenotyping approaches, image- and visual-based, in both populations, with values of 0.91 and 0.93 for Popn-I and Popn-II, respectively (Fig 1; Supplementary Fig 2). Broad phenotypic variation in disease severity was noted across both populations (Supplementary Fig 3). In accordance with the image-based approach, the average disease severity varied between 16.32% and 88% for Popn-I and between 14.5% and 97.2% for Popn-II. Likewise, through visual observation, the average disease severity ranged from 6.7% to 93.3% for Popn-I and from 9.0% to 98.0% for Popn-II.

The phenotypic data was also binarized as “0” for more tolerant genotypes and “1” for more susceptible genotypes (Fig 1B). After binarization, there were 8 and 10 individuals in the “0” category within Popn-I for the image and visual approaches, respectively. In Popn-II there were 12 individuals in the “0” category for both approaches. The heritability estimates for anthracnose resistance trait in SHB blueberries, in Popn-I was about 0.47, whereas for Popn-II, it was 0.5 (Supplementary Fig 4), indicating a moderate level of heritability for this trait. The correlations in disease severity ratings in those two populations were computed using common genotypes as checks, which showed values of 0.40 and 0.53 for image and visual assessments, respectively.

### Genome-wide association analysis

A total of 38,379 SNPs, distributed across the 12 haploid blueberry chromosome-scaled scaffolds, were independently tested for association with anthracnose resistance trait employing a GWAS approach (Supplementary Fig 5, 6). For the continuous scale, in the combined populations (Comb_popn), no associations were found despite employing both visual and image approaches (Fig 2A, Supplementary Fig 7). While we identified significant associations in three chromosomes (1,2 and 11) using the visual approach within Popn-I, no associations were detected using the image approach (Supplementary Fig 8A, 9). Similarly, within Popn-II both visual and image approaches did not detect any significant associations (Supplementary Fig 8B, 10).

**Fig 2.**
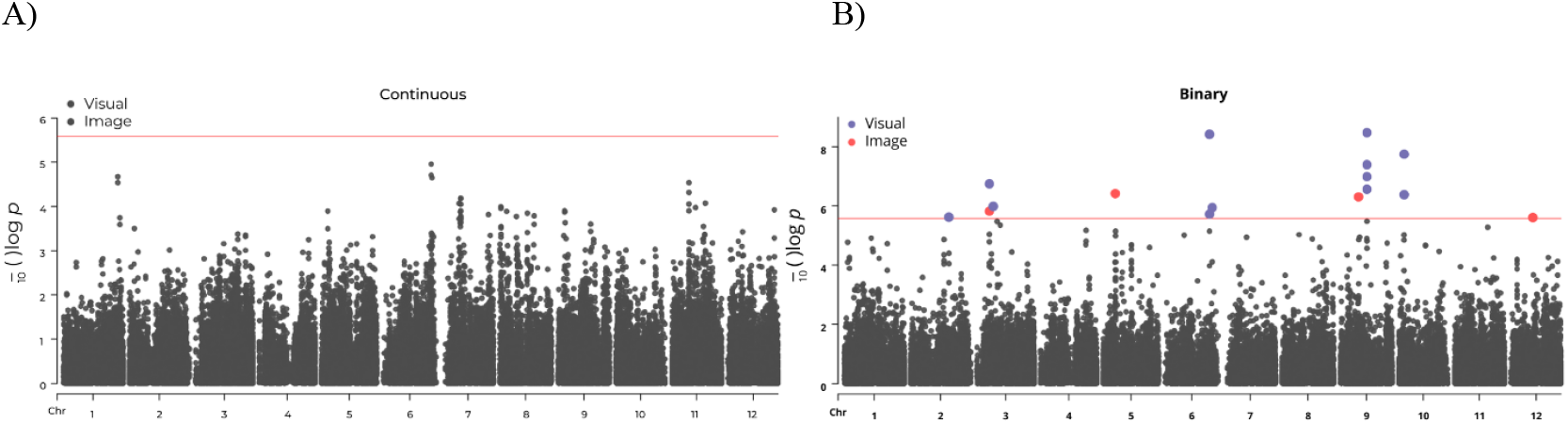
Manhattan plots for anthracnose resistance in the SHB population (Comb_Popn) using two distinct phenotyping methods: the image-based approach and the visual approach based on the A) continuous scale B) binarized scale. A linear mixed model with corrections for population structure and cryptic relatedness was used to compute the *p*-values. Adjusted Bonferroni correction, considering a genome-wide significance level of 0.05 (red line), was used to establish a *p*-value detection threshold for statistical significance.

With the hypothesis that the unaccounted defoliation discussed earlier may have limited our ability to detect significant associations when the continuous disease severity scale was employed, we further explored the results from the binarized dataset only. With the binarized dataset, varying numbers of significant SNPs were detected across different populations (Table 1). In the Comb_popn, significant associations were identified on chromosomes 3, 5, 9 and 12 using the image dataset, while the visual dataset revealed associations on chromosomes 2, 3, 6, 9 and 10 (Fig 2B, Table 1, Supplementary Fig 11). When each population was further analyzed independently, we uncovered significant associations on chromosomes 8, 9 and 11 in Popn-I (Fig 3A, Table 1, Supplementary Fig 12), that were consistently detected by both phenotyping approaches. However, an additional association on chromosome 3 was observed when employing the image approach. Notably, although associations were detected on the same genomic region, they were not at the same location (SNP position). In Popn-II (Fig 3B, Table 1, Supplementary Fig 13), we detected the same associated SNPs on chromosomes 5 and 6 across both phenotyping approaches. However, other non-overlapping associations were detected with image approach (on chromosomes 3, 8, 10 and 12) and with visual approach (on chromosome 2 and 12). Intriguingly, markers in chromosome 10 from Comb_popn (visual dataset) and from Popn-II (image dataset) were 23kb apart, pointing to a close genomic region. A significant marker in chromosome 12 detected in Comb_popn (image dataset) was also identified in Popn-II (visual dataset). This observation underscores the possibility of shared genetic loci influencing anthracnose resistance across populations. However, the major effect QTL observed in Popn-I on chromosome 8 was not consistent across populations, suggesting potential population- and environment-specific genetic determinants influencing anthracnose resistance.

**Table 1:**
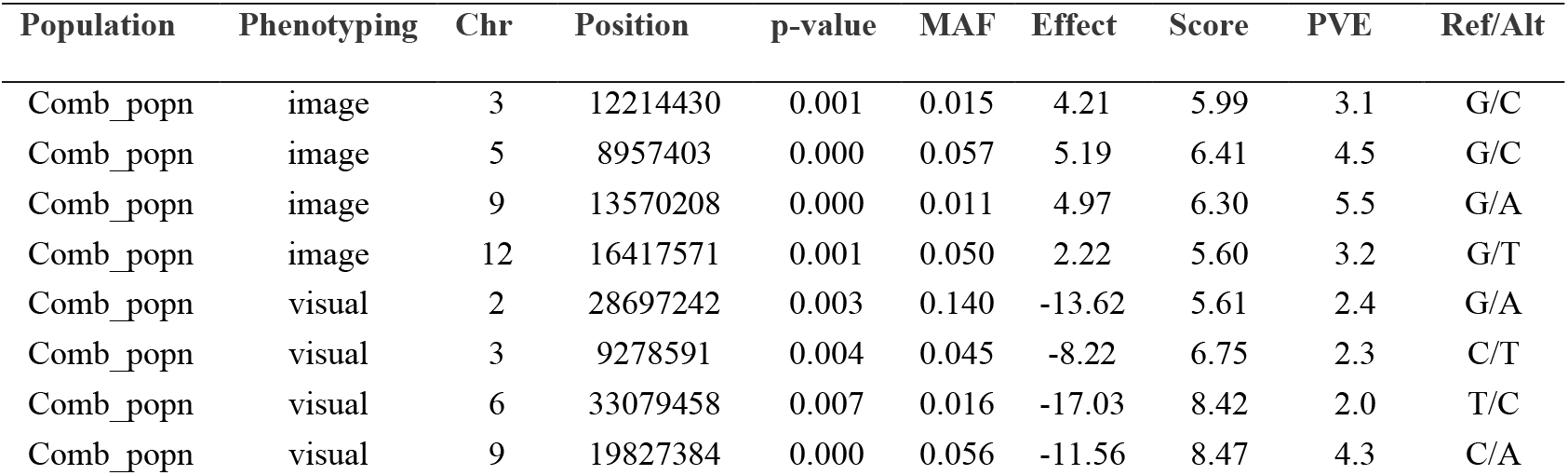
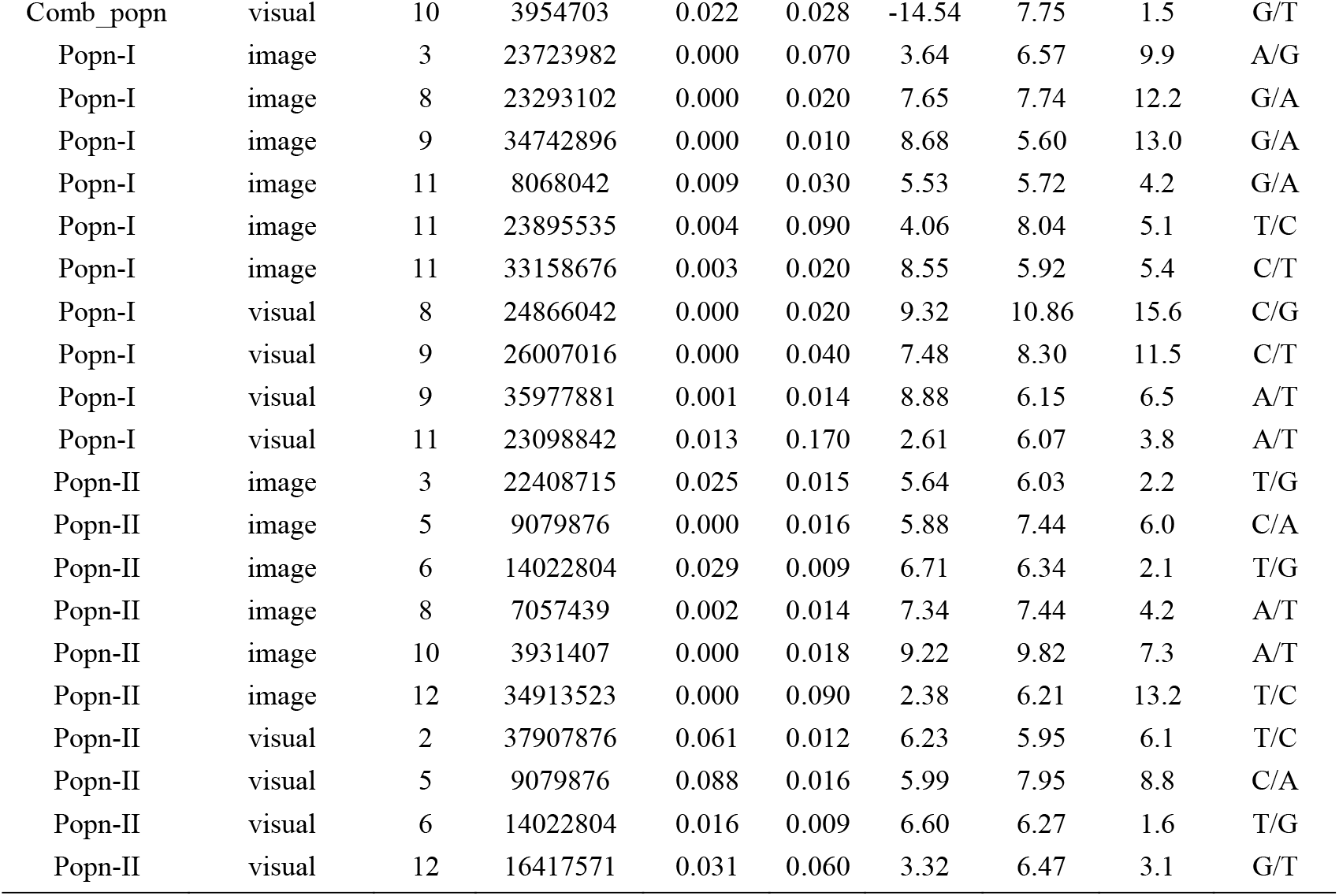
Significant SNPs associated to anthracnose based on binarized scale in SHB populations: in the combined population (Comb_popn) and in two independent populations: Popn-I and Popn-II, using two different phenotyping approaches: image- and visual-based. The corresponding SNP location, *p*-value, minor allele frequency (MAF), effect, score (i.e. -log_10_(*p*-value)), percentage of variance explained (PVE), and reference (Ref) and alternative (Alt) allele are reported.

**Fig 3.**
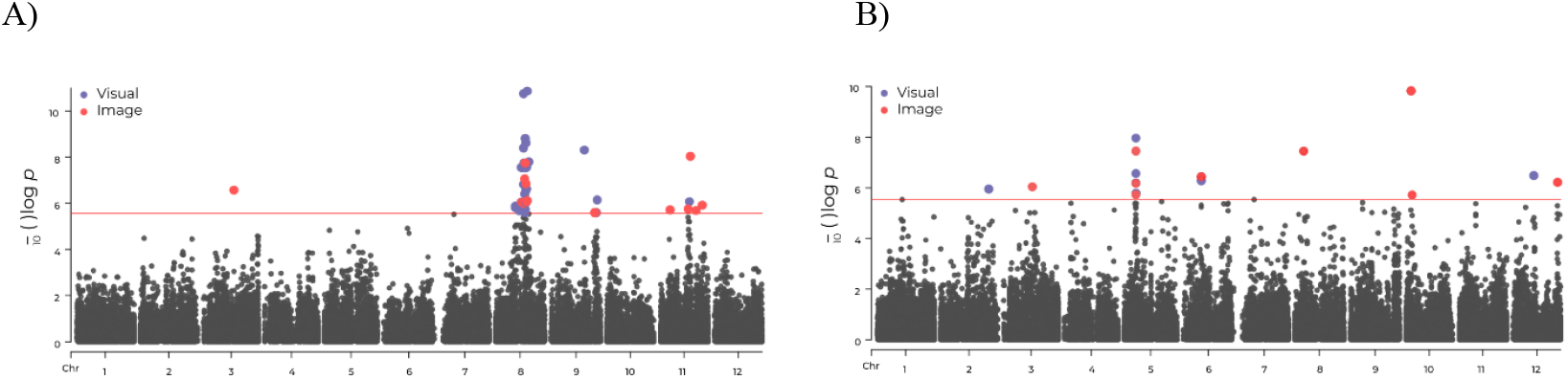
Manhattan plots for anthracnose resistance in the SHB population, (A) Popn-I (B) Popn-II, based on the binarized scale using two distinct phenotyping methods: the image-based approach and the visual approach. A linear mixed model with corrections for population structure and cryptic relatedness was used to compute the *p*-values. Adjusted Bonferroni correction, considering a genome-wide significance level of 0.05 (red line), was used to establish a *p*-value detection threshold for statistical significance.

### Candidate gene mining

Putative protein-coding genes within a 100kb window upstream and downstream of significant SNPs obtained in binary scale were retrieved from the reference genome. Candidate genes related to different layers of resistance mechanism are presented in Table 2. Genes encoding for pentatricopeptide repeat-containing protein, E3 ubiquitin protein ligase, and serine/threonine protein kinase were recurrently identified flanking significant SNPs detected across all three population scenarios, utilizing either the image or visual datasets (Table 2, Supplementary Table 1). The functional relevance of these genes is considered further in the discussion.

**Table 2:**
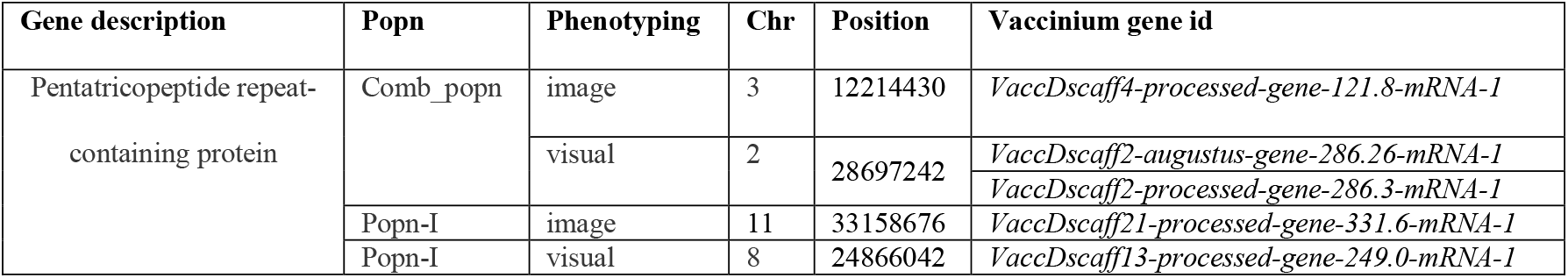
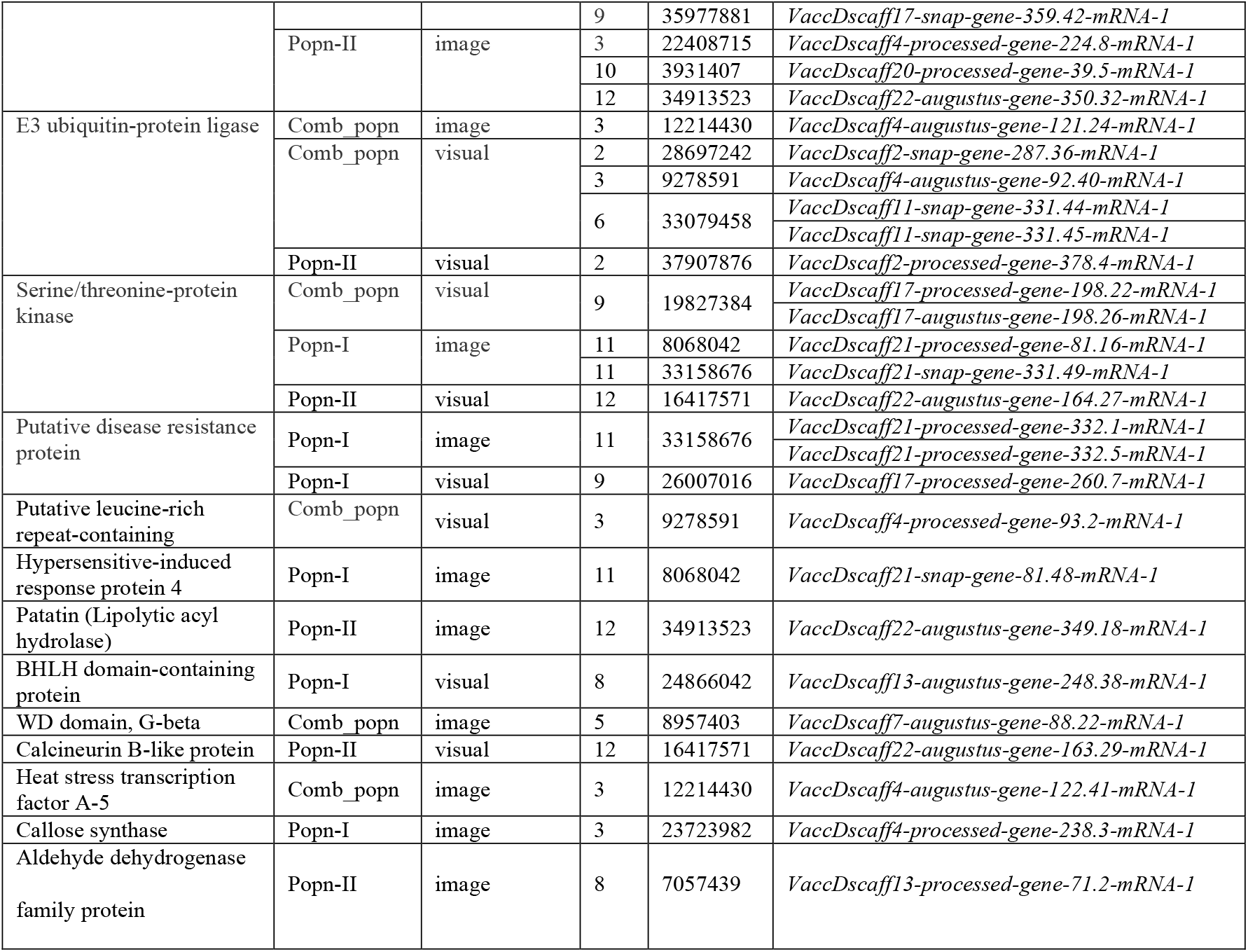
Selected candidate genes related to anthracnose resistance that are within ±100 kb of significant SNPs detected using binary scale.

## Discussion

Developing anthracnose-resistant blueberry cultivars relies on identifying resistance sources and understanding the genetic basis underlying the severity variability in the breeding population. Despite its importance, the genetic basis of anthracnose resistance caused by *C. gloeosporioides* remains understudied in blueberry. To address this gap, we conducted a GWAS using SHB blueberry breeding populations to elucidate the genetic and phenotypic variability and identify molecular markers and candidate genes associated with anthracnose resistance.

We inoculated detached stem branches of genotypes from two independent SHB blueberry populations with *C. gloeosporioides* and assessed the disease on the 35^th^ day post-inoculation with two phenotyping approaches: image- and visual-based. In our study, we observed instances of defoliation prior to disease severity rating in certain genotypes exhibiting moderate to severe symptoms of anthracnose, whereas leaf drop was not observed in genotypes with lower symptom levels. Defoliation may have removed leaves with severe symptoms disproportionately across genotypes with moderate to severe disease, potentially increasing variability in disease severity assessments, particularly for genotypes other than those exhibiting the most resistance. This unexpected experimental limitation may have impacted our ability to detect significant associations when the continuous disease severity scale was employed and would have impacted both visual and image analysis methods. This underscores the challenges associated with logistically capturing subtle genetic effects using continuous disease severity assessments on a large number of individuals.

To address this potential source of error, we decided to binarize the dataset, effectively looking only for genetic signal in the most resistant individuals as compared to all others combined. Binarization allowed us to focus on the most dramatic and potentially most useful differences and otherwise undetectable genetic associations. Drawing on this rationale, we employed a 5th percentile window based on the respective population’s average severity across three replicates to binarize our disease severity data. Individuals with an average disease severity value below the 5th percentile were categorized as 0, indicating minimal symptom development (i.e. resistant), while those above the 5th percentile were categorized as 1 (i.e. susceptible).

Our study revealed a high correlation between image and visual datasets, with the image approach (i.e. computer vision phenotyping) exhibiting higher sensitivity in detecting associations compared to the traditional visual approach across both studied populations with binarized data. This same approach of image analysis has demonstrated efficacy in evaluating other fruit quality traits, such as wax bloom in blueberry (Ferrão et al., *in press*). Moreover, image analysis techniques have proven advantageous in detecting new genetic loci for resistance against pathogens in various plant species, including tomato against *Ralstonia solanacearum*, Arabidopsis against *Sclerotinia sclerotiorum*, and wheat against *Zymoseptoria tritici* (Yates et al. 2019; Barbacci et al. 2020; Meline et al. 2023). Despite being more time-consuming than visual assessment in a study like ours, image analysis remains advantageous, particularly in scenarios when less-trained raters are involved in the scoring of these large populations. In blueberry breeding programs, accurate assessment of anthracnose severity is even more crucial as we observe its complex genetic architecture involving multiple genes (Osorio et al. 2014; Jacobs et al. 2020). So, integrating visual assessment with advanced image analysis techniques offers a promising solution to this challenge.

Leveraging these approaches can enhance precision and sensitivity of disease evaluation, minimize biases in traditional assessment methods, and capture more significant associations. In this regard, it is preferable to have false positives for further validation rather than missing potential targets.

We observed a moderate narrow-sense heritability estimate (∼0.5) for anthracnose resistance in SHB blueberries which closely aligned with the reported range of 0.63-0.73 for anthracnose fruit rot in northern highbush blueberries as per Miles and Hancock (2022) . This observation underscores the potential for breeding initiatives to effectively exploit genetic variation and enhance resistance against anthracnose, thereby contributing to the development of blueberry cultivars with improved disease resistance and thus a more sustainable production.

In our study, complete resistance to anthracnose was not observed among any genotype; however, high tolerance was observed in both populations. Moreover, the varying degree of response to anthracnose observed among different individuals underscores the presence of genetic variability within our population. Our GWAS analysis revealed the presence of significant associations across several genomic regions spanning different chromosomes. The presence of multiple minor additive genes contributing to a small portion of the phenotypic variance in blueberries is consistent to a complex and quantitative nature of anthracnose resistance as observed in other species like pepper, tea, cassava, sorghum, and beans (Oh et al. 2003; Boonchanawiwat et al. 2016; Perseguini et al. 2016; Mishra et al. 2017; Prom et al. 2019; Ro et al. 2023). While our study primarily focused on identifying genetic factors associated with resistance in leaf and stem tissues, potentially differing from those relevant in fruits, it’s imperative to acknowledge the experimental limitations in capturing broader field dynamics. There is a good possibility that the artificial inoculation method employed, characterized by high inoculum density (Jacobs et al. 2020) and close tissue proximity, may have overwhelmed many other minor resistance mechanisms. Therefore, while our findings provide valuable insights into genetic resistance mechanisms, they should be interpreted in the context of these experimental constraints.

In our observations certain chromosomes consistently showed significant associations across different populations. For instance, chromosome 9 had significant SNPs in both our study and a previous one by Jacobs et al. (2023) under scaffold 17, while chromosome 11 exhibited similar patterns within our study. However, it’s noteworthy that the location of significant SNPs in these chromosomes did not align with each other. One explanation for these differences in alignment is that many of the genes identified in this context are anticipated to play roles in diverse facets of host defense and are not predominant resistance genes. Despite this inconsistency, markers in chromosome 10 and 12 remained consistent in Popn-II and Comb_popn, indicating their potential as targets for marker-assisted selection.

Based on the significant genomic regions detected via GWAS, candidate genes putatively involved with defense pathways could be highlighted. While our study does not directly elucidate the molecular mechanisms and causal mutations underlying resistance, it provides potential candidates that could be targeted for further functional validation. The candidate genes identified encompass a range of functions, including pathogen recognition (such as resistance genes and resistance gene analogs [RGAs]) and general plant defense mechanisms and signaling. We reflect on the multifaceted nature of defense mechanisms and discuss genes associated with distinct defense layers.

Fundamental to plant defense is the process of pathogen recognition. Our study pinpointed RGA3, encoding a putative disease resistance protein, and a gene linked to a putative disease resistance RPP13-like protein. RGA proteins, encompassing nucleotide binding site leucine rich repeat (NBS-LRR) and transmembrane leucine rich repeat (TM-LRR) types, play vital roles in pathogen recognition and effector triggered immunity (ETI) response (Sekhwal et al. 2015). Previous research has highlighted *RPP13*’s association with *C. acutatum* and *C. gloeosporioides* resistance on grapes (Jang et al. 2020), and with *Peronospora parasitica* resistance on *Arabidopsis* (Bittner-Eddy et al. 2000). Additionally, we also detected another gene linked to pathogen detection, containing leucine-rich repeats (LRRs), essential for plant immune responses and defense against pathogens including *Colletotrichum* (Padmanabhan et al. 2009; Li et al. 2013; Jang et al. 2020; da Silva Dambroz et al. 2023).

Restricting the entry of pathogens and detoxification of reactive oxygen species are other vital aspects of plant defense. Plant cell wall is the primary target site of infection by the pathogen and so strengthening cell wall integrity to restrict pathogen growth is crucial. We identified a candidate gene encoding callose synthase, essential for synthesizing callose, a key component strengthening cell walls against pathogen intrusion (Zhang et al. 2023). Our research also detected a candidate pectinesterase-encoding gene crucial for plant defense, demonstrated across various species including banana, wheat, citrus and Arabidopsis, in response to diverse pathogens (Ma et al. 2013; L’Enfant et al. 2015; Zega and D’Ovidio 2016). Pectinesterases regulate defense against pathogens by de-esterifying pectin in cell walls, thus controlling access to pathogenic enzymes (Fan et al. 2017). Moreover, a gene encoding a hypersensitive-induced response protein was also identified, which plays a pivotal role in triggering rapid localized cell death at the site of pathogen invasion, effectively limiting pathogen spread including *Colletotrichum* and enhancing plant resistance (Biruma et al. 2012; Wang et al. 2018). Additionally, we observed a potential involvement of the heat stress transcription factor A-5 gene in regulating cell death induced by invasion of the pathogen or reactive oxygen species, as suggested by previous research (Baniwal et al. 2007). During stressful conditions, plants experience a significant increase in reactive oxygen species production across various cellular compartments, including mitochondria, chloroplasts, the endoplasmic reticulum, apoplast, peroxisomes, plasma membrane and cell walls (Kaur et al. 2022). So, reducing oxidative stress and limiting tissue damage from pathogens through detoxification is an important defense mechanism. Recent studies underscore vital role of pentatricopeptide repeat-containing proteins in mediating plant defense responses against *Colletotrichum* species in pepper (Ro et al. 2023) and sorghum (Prom et al. 2019), by modulating reactive oxygen species homeostasis within mitochondria and enhancing plant resilience (Geddy and Brown 2007; Laluk et al. 2011). In line with this, we also detected pentatricopeptide repeat-containing protein coding genes as potential candidates across multiple genomic regions. Another candidate gene, coding for an aldehyde dehydrogenase 1 (*ALDH*) family protein, was implicated in resistance mechanisms in tea and pepper fruit against *C. camelliae* and *C. gloeosporioides*, respectively (Oh et al. 2003; Wang et al. 2016). *ALDH* genes are reported to detoxify aldehydes like acetaldehyde, involved in stress signaling and disease resistance responses (Dolferus et al. 1994; Tadege et al. 1999). Our study speculated another gene encoding a BHLH domain-containing protein to be linked to anthracnose resistance, with evidence from grapevines suggesting its role in fungal-induced proanthocyanidins biosynthesis against *C. gloeosporioides* (Yu et al. 2023).

Signaling cascades is a critical layer of defense and coordinates plant responses to pathogen attacks and enhances overall immunity. Our GWAS study unveiled several genes involved in defense signaling cascades, including serine/threonine protein kinases and serine/threonine protein phosphatase. Previous research has demonstrated their involvement in resistance against *C. gloeosporioides* in walnuts (An and Yang 2014) and *C. lindemuthianum* in beans (Chen et al. 2017) and against other pathogens across diverse plant species (Martin et al. 1993; Song et al. 1995; Salmeron et al. 1996; Warren et al. 1998). Additionally, a gene encoding a WD domain, G-beta repeat-containing protein within family WD40, was identified that may also facilitate protein-protein interactions, and signaling pathways linked to defense responses (Villanueva et al. 2016). Likewise, another candidate gene encoding a calcineurin B-like protein has also been reported to play a vital role in plant defense by regulating calcium signaling pathways. Acting as a calcium sensor, it modulates the activity of downstream proteins involved in immune responses, as evidenced in other plant species (Luan 2009; Batistič and Kudla 2012).

Similarly, ubiquitin-mediated defense mechanisms are also involved in plant resilience against pathogens. Our study identified E3 ubiquitin ligase genes, and members of this family have been linked to resistance against *Colletotrichum* species in pepper (Ro et al. 2023) and against *Puccinia* species in wheat (Shahinnia et al. 2022).

These genes regulate innate immunity (You et al. 2016) and phytohormone signaling, enhancing antimicrobial defense and contributing to broad-spectrum disease resistance (Kelley 2018). Additionally, we found other genes within the ubiquitin pathway, coding for ubiquitin-conjugating enzyme E2 and RING-type E3 ubiquitin transferase.

Another crucial layer of plant defense lies in hormonal signaling modulation. Our study detected a gene encoding a Patatin-like protein that regulates the expression of jasmonic acid and ethylene biosynthesis genes, associated to resistance mechanisms in *Arabidopsis thaliana* and *Nicotiana attenuata* against fungal pathogens (La Camera et al. 2005; Cheng et al. 2019). Additionally, clusters of cytochrome p450 genes were also found in our study, and members of this family have been linked to resistance of pepper against *C. gloeosporioides* (Oh et al. 1999; Ko et al. 2005).

Although still speculative, our findings together with evidence from independent studies, particularly against *Colletotrichum* species, strengthen the hypothesis regarding the involvement of these candidate genes in resistance mechanisms. We acknowledge the possibility that other nearby genes and uncharacterized genes might be involved in the mechanism of resistance.

Altogether, the wide variability of severity symptoms across genotypes and the detection of several minor additive genes potentially involved in multiple layers of pathogen defense suggest that anthracnose resistance in SHB is polygenic and is quantitatively inherited. The potential candidate genes detected in this study likely confer non-race-specific resistance, since they are involved in signal transduction components and general biochemical defenses, such as cell wall reinforcement, production of reactive oxygen species, and induction of pathogenesis-related proteins (Miles et al. 2011; Jacobs et al. 2023). Developing and deploying cultivars with non-race-specific resistance could enhance durability compared to pyramiding major resistance genes (Michel et al. 2023). Thus, the next essential steps in our research involve assessing the usefulness of molecular markers for predicting phenotypes and validating candidate genes from our GWAS analyses. This is crucial for informing breeding strategies and ensuring the reliability of genetic associations, advancing our understanding of anthracnose resistance in SHB blueberries.

## Conclusion

This study aimed to elucidate the genetic basis of anthracnose resistance on blueberry leaves and stems. We established a screening protocol to assess anthracnose severity and conducted evaluations in two independent populations, consisting of advanced selections from the UF Blueberry Breeding Program. Additionally, we compared disease quantification using the computer vision phenomics with human vision. Then, through GWAS, we identified several genomic regions associated with anthracnose resistance against *C. gloeosporioides*. Our results underscored the quantitative inheritance of this trait, with multiple variants of minor effects contributing to resistance. These findings suggest a non-race-specific resistance mechanism, supported by the involvement of candidate genes in various defense pathways which can now be functionally validated and utilized in breeding programs. The screening protocol employed in this study can be integrated into the breeding pipeline to screen further selections against anthracnose, and the identified markers can serve as valuable genomic resources for expediting the development of anthracnose-resistant blueberry varieties.

## Acknowledgements

Financial support provided by the USDA National Institute of Food and Agriculture (NIFA) Award No. 2019-67013-29166. The authors would like to thank Dr. Norma Flor for her insights during the screening protocol development stage.

## Data availability

The authors affirm that all data necessary for confirming the conclusions of the article are present within the article, figures, tables and supplementary files. File S1 contains genotypic and File S2 contains phenotypic data.

## Contributions

PM conceived and supervised the study. PM, PH, and LG contributed to the study design. LG conducted the phenotyping, data analysis, and manuscript writing. TJ and BL developed the image-analysis pipeline. PA assisted with the data analysis. PH supervised the phenotyping process, while LF and CA oversaw the data analysis. JB and FE supervised the gene mining. All authors approved the final manuscript.

## Supplementary Data

Supplementary_Figs.doc and Supplementary_Table.xlsx

